# Warming undermines emergence success in a threatened alpine stonefly: a multi-trait perspective on vulnerability to climate change

**DOI:** 10.1101/2022.08.01.502337

**Authors:** Alisha A. Shah, Scott Hotaling, Anthony Lapsansky, Rachel L. Malison, Jackson H. Birrell, Tylor Keeley, J. Joseph Giersch, Lusha M. Tronstad, H. Arthur Woods

**Author notes:** **Correspondence:** Alisha A. Shah, W. K. Kellogg Biological Station, Department of Integrative Biology, Michigan State University, Hickory Corners, MI, 49060, USA; Scott Hotaling, Department of Watershed Sciences, Utah State University, Logan, UT, USA. **Author contributions:** Alisha Shah and Scott Hotaling designed the study with substantial input from Joe Giersch and Art Woods; Alisha Shah, Scott Hotaling, Joe Giersch, Jackson Birrell, Rachel Malison, Anthony Lapsansky and Tylor Keeley collected the data; Alisha Shah, Anthony Lapsansky, and Art Woods performed statistical analyses; Alisha Shah and Scott Hotaling wrote the manuscript. All authors contributed substantially to revisions and gave final approval for submission. **Data availability:** All data used in this study will be made available on Dryad, and all R code will be accessible via GitHub upon publication.

## Abstract

Species vulnerability to global warming is often assessed using short-term metrics such as the critical thermal maximum (CTmax), which represents an organism’s ability to survive extreme heat. However, an understanding of the long-term effects of sub-lethal warming is an essential link to fitness in the wild, and these effects are not adequately captured by metrics like CTmax. The meltwater stonefly, *Lednia tumana*, is endemic to high-elevation streams of Glacier National Park, MT, USA, and has long been considered acutely vulnerable to climate change-associated stream warming. In 2019, it was listed as Threatened under the U.S. Endangered Species Act. This presumed vulnerability to warming was challenged by a recent study showing that nymphs can withstand short-term exposure to temperatures as high as ~27 °C. But how this short-term tolerance relates to chronic, long-term warming has remained unclear. By measuring fitness-related traits at several ecologically relevant temperatures over several weeks, we show that *L. tumana* cannot complete its life-cycle at temperatures well below the CTmax values measured for its nymphs. Although warmer temperatures maximized growth rates, they appear to have a detrimental impact on other key traits (survival, emergence success, and wing development), thus extending our understanding of *L. tumana’s* vulnerability to climate change. Our results call into question the use of CTmax as a measure of thermal sensitivity, while highlighting the power and complexity of multi-trait approaches to assessing climate vulnerability.

## Introduction

Climate change is occurring more rapidly at high elevations than almost anywhere else on Earth (Pepin et al., 2015). This change has led to massive recessions of mountain glaciers and snowfields (Hugonnet et al., 2021) and contemporary warming of headwater streams (Niedrist & Füreder, 2020). In the future, rising stream temperatures will force mountain stream biota to either tolerate their new thermal regimes or be locally extirpated (Birrell et al., 2020; Hotaling, Finn, Giersch, Weisrock, & Jacobsen, 2017; Jacobsen, 2020), putting many species at risk of extinction (e.g., Giersch, Hotaling, Kovach, Jones, & Muhlfeld, 2017). In North America, two alpine stoneflies have been listed under the U.S. Endangered Species Act due to climate change-induced habitat loss (*Lednia tumana* and *Zapada glacier*, US Fish and Wildlife Service, 2019). However, the vulnerability of alpine stream insects has rarely been measured, particularly beyond short-term tests of thermal tolerance.

Vulnerability to climate change is often assessed using short-term metrics such as the critical thermal limits—CT_MAX_ and CT_MIN_—which represent, respectively, the high and low temperature limits of an organism and are marked by a loss of locomotor capacity (Angilletta, 2009; Chown, Jumbam, Sørensen, & Terblanche, 2009; Lutterschmidt & Hutchison, 1997; Sinclair et al., 2016). Although easy to measure, and powerful in a comparative framework (Addo-Bediako, Chown, & Gaston, 2000; Dallas & Rivers-Moore, 2012; Shah et al., 2017; Sunday, Bates, & Dulvy, 2012), estimates of CT_MAX_ and CT_MIN_ are difficult to link directly to fitness in the wild (Huey & Berrigan, 2001). In general, single traits represent only components of fitness, and each trait may vary to different degrees across a thermal range (Kingsolver & Woods, 1997; Shah, Bacmeister, Rubalcaba, & Ghalambor, 2020a). For example, in a comparison of temperate and tropical mayflies, thermal breadth (i.e., the difference between CT_MAX_ and CT_MIN_) was narrower for tropical mayflies indicating that they are more vulnerable to warming than temperate mayflies (Shah et al., 2017). However, for the same groups, swimming performance did *not* vary significantly suggesting that from a performance standpoint, they are equally sensitive to warming (Shah et al., 2020a). These seemingly opposing conclusions about vulnerability can be addressed and clarified by measuring multiple fitness-related traits (Kellermann & Heerwaarden, 2019; Pearson et al., 2014).

Critical thermal limits alone may provide only a partial understanding of vulnerability because they are typically measured over just a few hours and represent thermal ‘end points’ (Hochachka & Somero, 2002). However, the absolute temperatures that animals can tolerate are not fixed, and they decrease with duration of exposure (Rezende, Castañeda, & Santos, 2014). Critical thermal limits also do not reflect organismal responses to the slower, chronic nature of climate warming, where performance is not dictated by survival alone, but also by growth, development, and reproduction (Verberk, Durance, Vaughan, & Ormerod, 2016). For most wild organisms, little is known of the long-term effects of modestly higher temperatures on fitness, and whether warming will be a net cost or benefit. For instance, some ectotherms may grow and develop faster in warmer conditions (Deutsch et al., 2008), but fast growth itself may trade off with other important traits (e.g., starvation resistance, development, or immune function; Arendt, 1997) and may produce less robust or even abnormal adults (Arnott, Chiba, & Conover, 2006; Ficetola & De Bernardi, 2006). When reared at warmer temperatures, for example, fruit fly larvae grow faster but fly more slowly as adults due to under-developed wing muscles (Fraimout et al., 2018).

Assessing vulnerability to climate warming from single traits is particularly challenging for organisms with complex lifecycles, such as aquatic insects whose life histories bridge aquatic and terrestrial environments. Aquatic juveniles typically experience less thermal variation in water, but they also have fewer opportunities to escape from suboptimal temperatures. By contrast, adults experience greater thermal variation as well as more microclimatic options for thermoregulation (Shah, Dillon, Hotaling, & Woods, 2020b; Woods, Dillon, & Pincebourde, 2015). These environmental differences likely drive divergent physiological tolerances between immature and adult stages. Nevertheless, conditions that juveniles experience can carry over beyond the developmental environment and influence adult traits (Bonte, Travis, Clercq, Zwertvaegher, & Lens, 2008; McCauley, Hammond, & Mabry, 2018). Because most aquatic insects spend a large proportion of their lives in water before emerging as winged adults, changes in both aquatic and terrestrial environments can impact their responses to climate change.

Stoneflies in the genus *Lednia* (Family: Nemouridae) are endemic to alpine habitats of western North America (Green et al., 2022). *Lednia tumana* occur in alpine streams of Glacier National Park, USA, and the surrounding mountains including many glacier-fed streams. Given their close association with ice and snow, *Lednia* species are presumed to be highly vulnerable to climate warming (Green et al., 2022), but this prediction is based on correlations between abundances and stream temperatures (Giersch et al., 2017; Green et al., 2022) and lacks a mechanistic explanation. Still, in 2019, *L. tumana* was listed under the U.S. Endangered Species Act due to climate change-induced habitat loss (U.S. Fish & Wildlife Service, 2019). Recent work has challenged the basic tenet of vulnerability for *L. tumana* by showing that two species in the genus—*L. tumana* and *L. tetonica*—can withstand short-term exposure to temperatures that are remarkably high compared to the in-stream temperatures they naturally experience (up to an average of 27.4 °C; Hotaling et al., 2020). Further, *L. tumana* and other members of its “cold water” community persist in alpine basins that have not been glaciated in ∼170 years (Muhlfeld et al., 2020). Thus, there is a disconnect between where the species occurs, its presumed vulnerability, and the limited data on its physiological tolerances.

Here, we attempted to resolve this debate by experimentally assessing the effects of prolonged exposure to ecologically relevant temperatures on multiple development and performance-based traits for *L. tumana* nymphs and adults. Specifically, we addressed three questions: (1) What are the highest temperatures *L. tumana* nymphs can withstand over multi-week timescales? (2) How do different traits vary in their responses to this prolonged warming? And, (3) how does long-term exposure to warmer temperatures during aquatic development affect the performance of terrestrial adults? Beyond new understanding of climate risks for a threatened species, our results provide the first multi-trait perspective on climate vulnerability for any alpine stream insect. This is particularly relevant in light of global threats to alpine stream invertebrates (Birrell et al., 2020; Brown, Hannah, & Milner, 2007; Jacobsen, Milner, Brown, & Dangles, 2012; Leys, Keller, Robinson, & Räsänen, 2017), and our results have direct implications for their persistence under climate change.

## Materials and Methods

### Insect collection and incubation

We collected mid-to-late instar *Lednia tumana* nymphs (*N* = 419) from five streams in the alpine of Glacier National Park, MT, USA in mid-July 2019 (Fig. 1; Table 1). In each stream, we deployed temperature loggers (HOBO Water Temperature Pro v2 data; Onset Computer Corporation) secured to rocks with coated steel cable. Nymphs were transferred from the stream in Whirlpak bags filled with stream water to the University of Montana in Missoula, MT. At the time of collection, we also scraped rocks to collect algae and biofilm (hereafter “biofilm slurry”), typical food sources for *L. tumana* and many other nemourids (Karen Jorgensen, *unpublished data*; Lieske & Zwick, 2007). The biofilm slurries were collected in Lunch Creek, a representative stream with a very high density of *L. tumana*, and used to feed nymphs during the experiment.

**Table 1:** Sampling sites and locations included in this study. Site temperature average (T_AVG_), minima (T_MIN_), and maxima (T_MAX_) are in degrees Celsius. At each site, temperature was measured during a ∼5-day period in the summer (13 – 17 August 2019).

| Site | Elevation(m) | Latitude, Longitude | $T_{\text{AVG}}$ | $T_{\text{MIN}}$ | $T_{\text{MAX}}$ | $N_{\text{SAMPLED}}$ |
| --- | --- | --- | --- | --- | --- | --- |
| Siyeh Bend | 1830 | 48.7002°, -113.6701° | 7.7 | 2.6 | 14.7 | 97 |
| Sexton Glacier | 1893 | 48.7003°, -113.6250° | 3.0 | 1.6 | 5.3 | 110 |
| Lunch Creek | 2101 | 48.7049°, -113.7041° | 6.5 | 4.1 | 10.8 | 105 |
| Going-to-the-Sun | 1746 | 48.6944°, -113.6201° | 4.4 | 2.9 | 6.3 | 54 |
| Reynolds Spring | 2085 | 48.6818°, -113.7322° | 7.1 | 2.8 | 14.5 | 53 |

**Figure 1:**
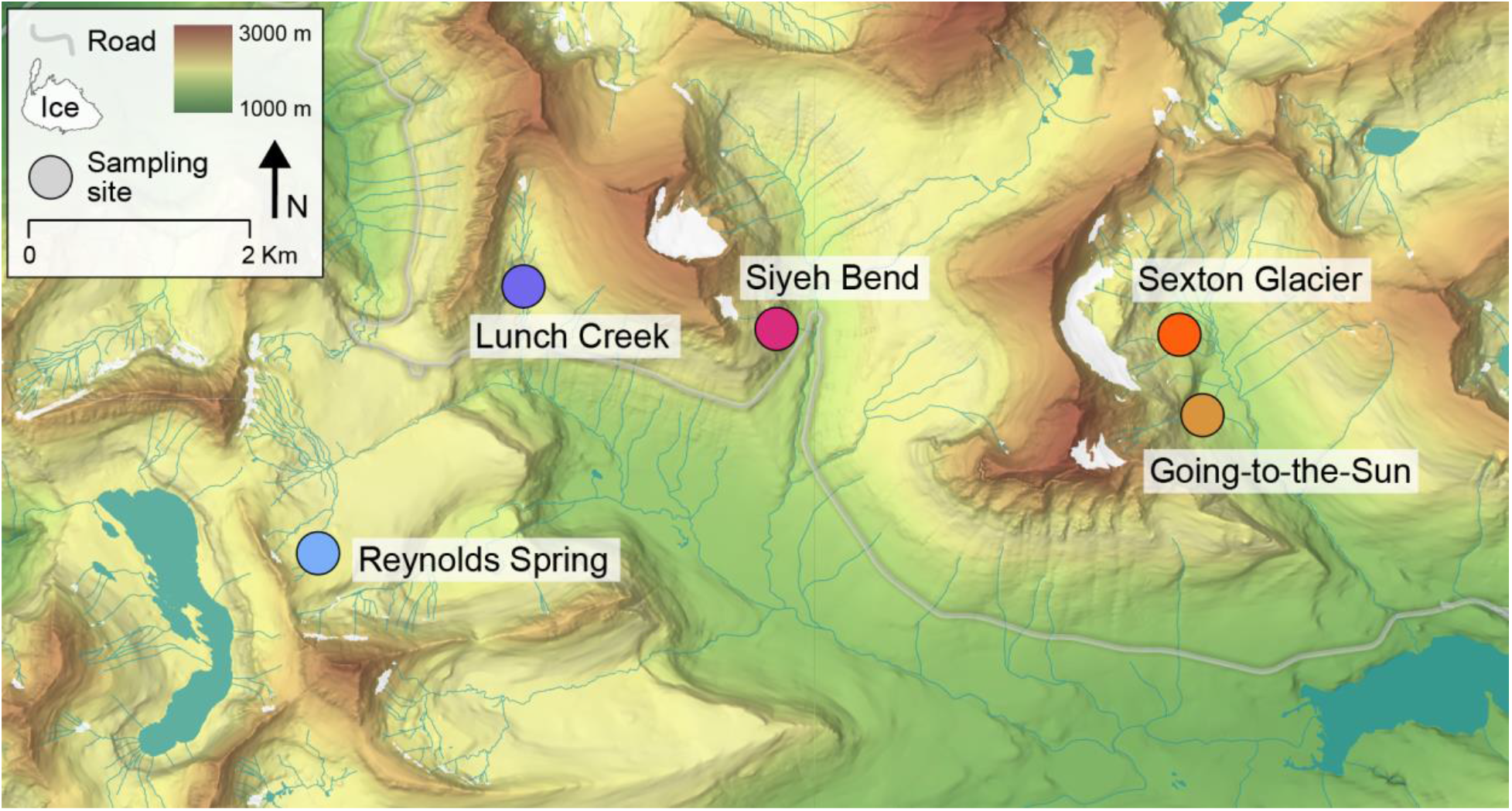
A map of our study sites in Glacier National Park, MT, USA where *Lednia tumana* nymphs were collected.

In the laboratory, nymphs were transferred to incubators (Percival Scientific, LT-36VLC8, I-66LLC8, I-36LLC8, I-66LLC8) that were initially held at 7 °C to match the water temperature in the Whirlpak bags after transport. Over ∼5 hours, incubators gradually reached fixed test temperatures of 1, 4, 7, 13, 21 °C. We also included a variable treatment (T_MAX_ = 10 °C, T_MIN_ = 4 °C, average = 7 °C) to mimic the summer temperature regime of a thermally variable stream, Lunch Creek, included in our study. For the variable treatment, the incubator rose to the maximum temperature (10 °C) over 3 hours (0700 – 1000 hours), remain stable for 9 hours (1000 – 1900 hours), then decrease to the minimum (4ºC) over 3 hours (1900 – 2200 hours), which was then held overnight.

For all treatments, light-dark cycles were set to 16:8 h L:D to approximate summertime in the northern Rockies. Each population × temperature treatment had two replicates of 7 individuals on average (max = 10, min = 3). Replicate groups of nymphs were placed in separate clear plastic containers with air stones for oxygenation and fed with 100 mL of biofilm slurry. We placed two ceramic tiles (2 cm diam.) in each plastic container to serve as substrate. Throughout the study, levels of food and water were checked once per day. The experiment lasted 31 days.

### Trait measurements and analyses

We measured growth, emergence success, survival, and adult locomotor performance in *L. tumana* across 6 temperature treatments (5 fixed temperatures and 1 variable regime). To obtain growth data, nymphs were photographed once per week from above (Olympus Tough TG-6) with a 2 cm scale placed in the field of view. To allow for assessment of emergence success, we placed plastic ladders, cut from embroidery mesh, in each container and stretched soft tule mesh over a wire frame to trap adults as they emerged. Each day the number of emergences and mortality were noted. Dead individuals were removed, and emerged adults were gently captured within one day of emergence and held at 7 °C for up to 2 days for locomotor performance measurements. All data analyses were conducted in R (R version 3.6.3).

### Growth

We measured body length (tip of head to tip of abdomen, not including cerci) from photos using the line tool in Image J. To obtain a more energetically-based response variable from body length, we converted length into dry mass using a length-mass regression:

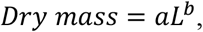

where, *a* and *b* are constants obtained from length-mass regressions of other nemourid species (Benke, Huryn, Smock, & Wallace, 1999), and *L* is body length. Because it was not possible to tell individuals apart within containers, we calculated a mean dry mass for each replicate container. We found that growth had ceased in some of the warmer treatments after two weeks. This is likely because nymphs had completed growth and were very close to emergence. We therefore restricted our growth analyses to only the first two weeks of the experiment. P.er-week growth was measured as the difference between mean mass between the first two weeks for each replicate divided by 2. Due to small sample size, we pooled replicates for each site × temperature for analyses. Then, using the R package *nls2*, we fit a thermal performance curve to the growth data using a Gaussian function (Angilletta, 2006):

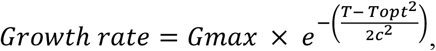

where, *Gmax* is the height of the peak, *T* is the sequence of temperature values over which growth was assessed, *Topt* is the position of the center of the peak, and *c* is the standard deviation of the curve. The starting parameter values were: *Gmax* = 0.07, *Topt* = 13, and *c* = 5.

### Survival

We analyzed survival using a logistic regression implemented as a generalized linear model (glm, family ‘binomial’) in R. For each individual, survival was coded as a binomial variable, i.e., as a 1 (survived) or a 0 (died). Emerged insects were also counted as having survived. We then assessed the fixed effects of temperature, site, and their interaction on survival and used the random effect of container to account for the grouping of individuals into different containers.

### Emergence

We assessed the effect of temperature on emergence using a linear model. We used proportion emerged as a response variable and the interaction term temperature x site as a predictor variable to test whether emergence differed across temperature for different sites.

### Adult locomotor performance and wing morphology

Flight is the primary mode by which adult aquatic insects swarm to find mates and disperse. In addition to flying in air, adult stoneflies also skim across the water surface to move across a stream. When surface skimming, stoneflies use their wings to generate aerodynamic thrust without leaving the surface of the water (Marden & Kramer, 1994). This form of locomotion is widespread across Plecoptera (Marden, O’Donnell, Thomas, & Bye, 2000; Thomas, Walsh, Wolf, McPheron, & Marden, 2000) and relatively well-studied because of its hypothesized role in the evolution of insect flight (Marden & Kramer, 1994; Will, Marden, & Kramer, 1995). We used the average sustained speed of surface skimming in *L. tumana* to test whether the temperature they experienced as juveniles affected their adult performance.

Adult insects were dropped from their plastic holding container onto water in a metal tray (30 × 40 × 2 cm deep) by inverting and tapping the container. Skimming was filmed from above using a leveled, tripod-mounted Fastcam Mini AX100 (Photron, Tokyo, Japan) recording at 2000 frames per second (see video in Supplementary Information). Each adult was dropped 3 to 4 times consecutively and filmed for 120 s each time.

Stoneflies were auto-tracked from videos using DeepLabCut (Mathis et al., 2018). To account for digitization error, we smoothed these data using MATLAB R2021a (Mathworks, Natick, MA) and the ‘smoothingspline’ function (The Mathworks, 2019). We plotted speed versus time for each skim and used the average speed of the longest non-zero and approximately horizontal portion of the plot (exhibited after initial acceleration but before final deceleration) to measure the average sustained skimming speed. Because laboratory temperature was difficult to precisely control, we recorded the temperature and adjusted the skimming speed to an estimated value at 22.5 °C by using the slope of a regression derived for the winter stonefly *Taeniopteryx burksi* (Marden & Kramer, 1994; Marden et al., 2000). This ‘adjusted skimming speed’ was used in downstream analyses. After skimming tests, adults were immediately preserved at −20 ºC. We assessed the effect of incubation temperature on adjusted skimming speed using a linear model with each individual’s mean adjusted skimming speed as a response variable and the interaction between temperature and site as a predictor variable in the model to test if temperature affected skimming performance differentially across sites.

We also measured wing length to use as a correlate of locomotor ability. We removed the fore- and rear wings from the right side of each preserved adult and photographed them (Nikon D7100) under a stereomicroscope (Nikon SMZ1500). Damaged wings were excluded from analysis. Wing length was measured as the distance between the upper corner of the wing base and the apex of the third radial (R3) wing vein near the wing tip using the Image J line tool. We pooled measurements within sites and used a linear model in R to test if developmental temperature affected fore- and rear wing length. We also measured wing area (see Supplemental Information, Fig. S1.), but because this metric was highly correlated with wing length (*r* _(31)_ = 0.93, *p* < 0.001), we focused on wing length in our analyses. To confirm that wing morphology is correlated with adult locomotor performance given our study design, we analyzed the relationship between each individual’s forewing and rear wing lengths and their sustained speed during surface skimming.

### Variable vs. static incubation treatment

To measure differences in trait sensitivity between individuals in the variable temperature treatment (min = 4 °C, max = 10 °C, mean = 7 °C) versus the most similar fixed temperature treatment (7 °C), we conducted Welch’s two-sample t-tests for each trait, i.e., growth, survival, emergence, and skimming performance.

## Result

### Growth

For all populations, growth rate was highest between weeks 1 and 2 of the experiment. Curve-fitting revealed that T_OPT_ lies at ∼13 ºC (Fig. 2a, Table 2).

**Table 2:**
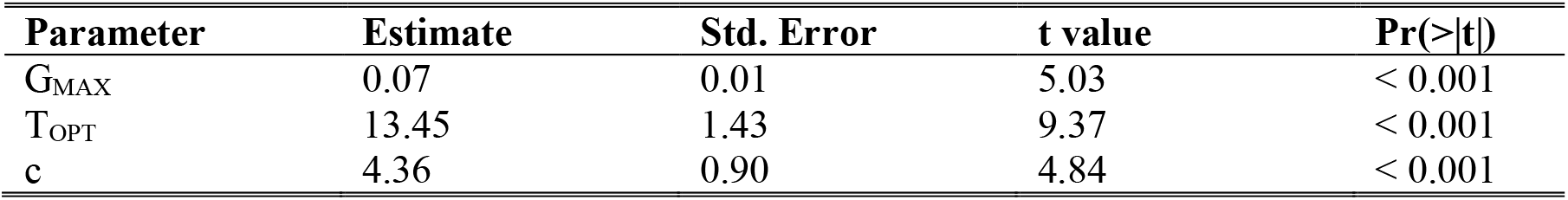
Parameter estimates of the thermal performance curve for growth in *Lednia tumana*.

**Figure 2:**
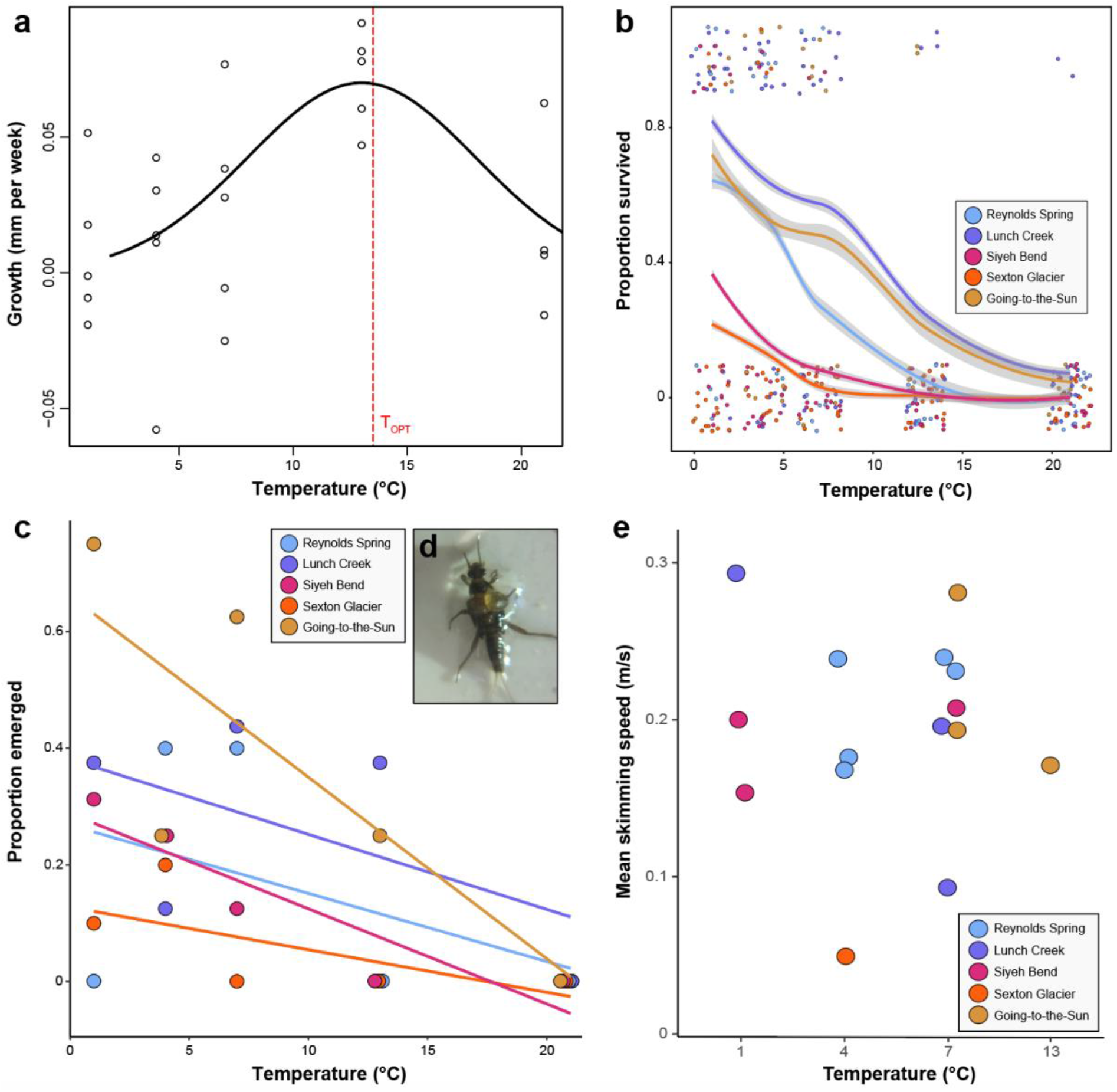
(a) A Gaussian function fitted to growth rates pooled across replicates for each site × temperature treatment. The dashed vertical line indicates T_OPT_ at 13.5 °C. (b) The proportion of individuals that survived in each temperature treatment. Survival declined with increasing temperature, and was notably low at 13 °C, the temperature at which growth rates were highest. For visualization, points are jittered on both the x- and y-axes. (c) The proportion of individuals that emerged. For all populations, emergence decreased with temperature and was exceptionally low at 13 and 21 °C. At 13 °C, several individuals appeared to be injured during emergence. Points are jittered along the x-axis wherever they overlapped. (d) An example of a nymph injured during emergence, with a partially split exoskeleton and a hemolymph bubble arising from its thorax. (e) Adult mean skimming speed across sites and temperature treatments. There was no variation in adult performance across sites or incubation temperatures.

### Survival

Survival declined with increasing temperature: individuals survived best at the coolest temperatures, 1 °C (55%), 4 °C (41.3%), 7 °C (33.5%) and the worst at the warmest temperatures, 13 °C (9.8%) and 21 °C (3.2%; Fig. 2b). Insects from Sexton Glacier and Siyeh Bend had consistently low survival across all temperatures. The logistic regression with temperature, site, and their interaction as predictors indicated that the main effects of temperature and site were significant (temperature: chi sq = 34.44, p < 0.001; site: chi sq = 32.37, p < 0.001) but that their interaction was not (chi sq 1.86, p = 0.76). Notably, survival for all sites was low in the 13 °C treatment (mean across populations = 9.8 %) where growth was highest.

### Emergence

The percentage of emerged individuals varied across populations but declined with increasing temperature (*F*_(1,15)_ = 11.36, *P* = 0.004). Specifically, more individuals emerged at the cooler temperatures: 1 °C (30.8%), 4 °C (24.5%), 7 °C (31.8%) and the fewer at the warmest temperatures, 13 °C (12.6%) and 21 °C (0.1%) (Fig. 2c). Only a few individuals emerged from the 13 °C treatment and none from the 21°C treatment (Fig. 2c). At 13 °C, some individuals exhibited dark wing pads (indicating they were ready for emergence), but with hemolymph leaking from their bodies (Fig. 2d). Hemolymph leakage may have been indicative of a heat-related injury and prevented completion of their final molt. There was no interaction between site and temperature treatment (*F*_(4,15)_ = 0.75; *P* = 0.57).

### Adult locomotor performance and wing morphology

Consistent with previous studies (e.g., Marden & Kramer, 1994) there was a significant relationship between trial-averaged skimming speed and both forewing length (*F*_(1, 12)_ = 8.20, *P* = 0.014, *R*^*2*^ = 0.41) and rear wing length (*F*_(1, 11)_ = 9.15, *P* = 0.012, *R*^*2*^ = 0.45; Fig. S2).

We then compared wing lengths between incubation treatments. Because few individuals with intact wings emerged from the 13 and 21 °C treatments, we could only compare wing characteristics among insects incubated at 1, 4, and 7 °C. Although there was a trend for longer fore- and rear wings in the 4 ºC incubation treatment (Fig. 3) we found no significant variation either in forewing length (*F*_(1, 31)_ = 0.64, *P* = 0.43; Fig. 3a) or rear wing length (*F*_(1, 27)_ = 0.16, *P* = 0.69; Fig 3b) among incubation temperatures. Skimming speed did not differ among adults emerging from different temperatures (*F*_(4, 12)_ = 0.61, *P* = 0.66; Fig. 2e) because there was no variation among wing lengths and incubation treatments, as shown above. There was also no significant interaction between site and incubation temperature on wing length (*F*_(4, 12)_ = 2.15, *P* = 0.18).

**Figure 3:**
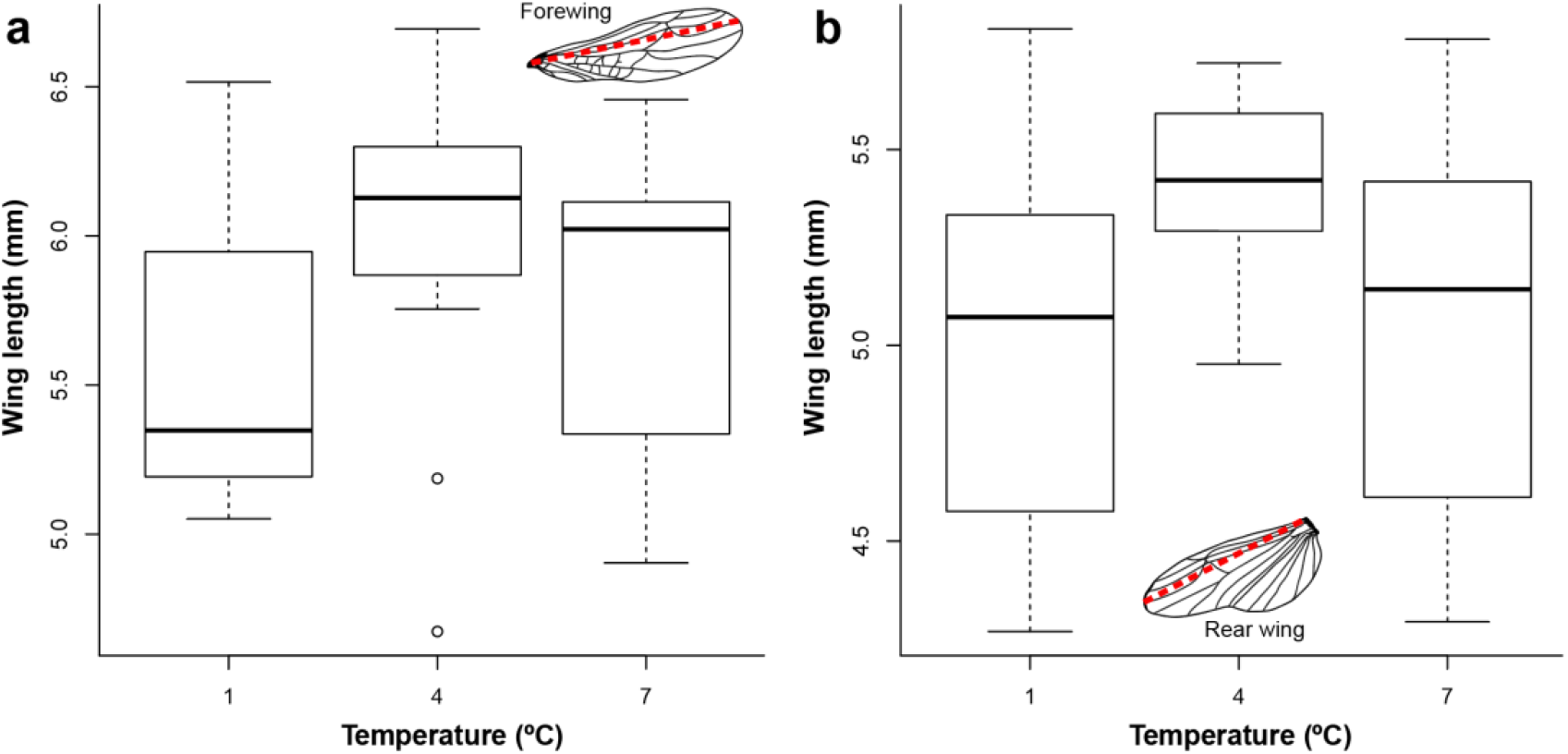
Adult wing length as a function of incubation temperature. There was no variation in wing length for different temperature treatments. However, for forewings (a) and rear wings (b), insects that emerged from the 4 °C treatment showed a trend for longer wings compared to those emerged from 1 °C and 7 °C. Forewings and rear wings were measured according to the dashed red line shown on representative wings in (a) and (b), respectively.

### Variable vs. fixed incubation treatment

Mean trait values were not different between insects reared in the variable temperature treatment (average = 7 °C) versus the fixed (7 °C) treatment (Table 3).

**Table 3.** Welch’s two-sample test statistics for difference between means in the variable (mean temperature = 7 ºC) versus fixed (7 ºC) treatments for all traits.

| **Trait** | **Df** | **t-statistic** | **P-value** |
| --- | --- | --- | --- |
| Growth | 12.59 | 2.01 | 0.07 |
| Survival | 4.85 | -0.50 | 0.64 |
| Emergence | 6.34 | 1.06 | 0.33 |
| Skimming speed | 3.05 | -0.68 | 0.55 |

## Discussion

Rising temperatures and shifting precipitation regimes are driving the recession of glaciers and permanent snowfields worldwide (Hugonnet et al., 2021, IPCC 2021) potentially endangering a multitude of species (Cauvy-Fraunié, Espinosa, Andino, Jacobsen, & Dangles, 2015; Stibal et al., 2020). For aquatic biodiversity, rising water temperature presents the most pressing risk associated with declines of the cryosphere (Hotaling et al., 2017; Niedrist & Füreder, 2020). However, the presumed risk of warming to aquatic species is largely derived from current distribution patterns, i.e., species restricted to meltwater are presumed to face greater risk of extirpation or extinction (e.g., Giersch et al., 2017). Although a useful starting point, assessing risk from distributions alone lacks an eco-physiological cause-and-effect context and may under-or overestimate the resilience of particular species to climate change. Gaining relevant eco-physiological insight through experimentation, however, can also be challenging, particularly for aquatic species endemic to remote regions, such as mountain tops, and whose life histories span aquatic and terrestrial habitats.

Short-term metrics of thermal tolerance (e.g., CT_MAX_) are easy to implement, and have therefore been widely used to identify species’ tolerance to warming (e.g., Bruno et al., 2018; Deutsch et al., 2008; Pinsky, Eikeset, McCauley, Payne, & Sunday, 2019). However, short-term metrics are sensitive to methodology, including temporal scale (Leiva, Calosi, & Verberk, 2019; Rezende et al., 2014) and do not address how species respond to chronically warm temperatures. Thus, there is a pressing need to consider responses that will likely be impacted at lower, sub-lethal temperatures.

Here, using multiple fitness-related traits, we evaluated the consequences of warming over ecologically relevant timescales for *Lednia tumana*, a threatened stonefly endemic to high-elevation streams in and around Glacier National Park, USA. The association between *L. tumana* and cold, often glacier-fed, meltwater has prompted investigations of its response to thermal stress (e.g., Hotaling et al., 2021, 2020; Treanor, Giersch, Kappenman, Muhlfeld, & Webb, 2013), but to date, this work has consisted of only short-term measurements (i.e., 1-3 hours) of tolerance to heat and cold stress. Overall, our results reveal that *L. tumana* is more sensitive to warming than short-term metrics have indicated. For example, even though CT_MAX_ indicates that *Lednia* can survive temperatures up to ∼27 °C, we show that lower temperatures (above 13 ºC) yield high rates of mortality at longer timescales. Moreover, thermal performance differed among traits; nymphs grew fastest at higher temperatures, whereas they transitioned into adulthood (emergence) best at much lower temperatures, with most individuals emerging between 1 and 7 °C. Thus, there is clear value in determining which traits are impacted most by warming to more accurately predict climate change outcomes. Generally, our results indicate that mountain streams that exceed 7 °C for prolonged periods in the summer may not support *L. tumana* populations.

### Temperature effects on growth, survival, and emergence

Despite low emergence rates above 7 °C, growth was maximized at higher temperatures (∼13 °C). For small ectotherms living in temperate and high-elevation regions, growth is often limited by low temperatures (Woods et al., 2022; Roitberg & Mangel, 2016). At warmer temperatures, life processes such as metabolic rate, food intake, and rates of enzyme mediated reactions, tend to peak at temperatures organisms experience in the wild (Angilletta & Dunham, 2003). Many temperate insects including *L. tumana*, perform best at warm temperatures (Frazier, Huey, & Berrigan, 2006), likely higher than those they experience in the wild. Assuming that stream productivity will increase with warming (Oleksy et al., 2020) and fuel faster growth (Miller, Clissold, Mayntz, & Simpson, 2009), high T_OPT_ for growth suggests that some *L. tumana* populations may benefit from marginally warmer streams, at least initially (Deutsch et al., 2008). However, additional research on important traits such as feeding and metabolic rates, as well as egg count of *L. tumana* will be essential to understanding the full effects of temperature on fitness.

In contrast to growth, other life-history traits in *L. tumana* showed greater sensitivity to temperature, which undermines conclusions about benefits and harm, and underscores the importance of investigating multiple traits. Survival declined steeply across populations with increasing temperature, and, emergence was low (∼10%) at the temperature that yielded the highest growth rate (13 °C). At this temperature, many emerging adults died before exiting the water, often with hemolymph leaking from their thoraxes. Adults from cooler treatments did not show similar injuries. High growth but low emergence at 13 ºC may reflect an underlying trade-off driven by a lack of adequate energy supply to fuel a complete emergence in warm water (Nash, Antiqueira, Romero, Omena, & Kratina, 2021). This increased energy requirement for molting may compound an already rising demand for oxygen at warmer temperatures (Dallas & Ross-Gillespie, 2015; Verberk et al., 2016).

### Temperature effects on adult traits

Temperature is a well-known determinant of ectotherm size at adulthood, and particularly, lower temperatures during development often result in larger adult size (French, Feast, & Partridge, 1998; Ray, 1960). For winged ectotherms, temperature can also affect the morphology and, ultimately, flight performance (Frazier, Harrison, Kirkton, & Roberts, 2008; Verberk et al., 2021). We examined the effects of developmental temperature on wing traits and skimming performance. Although adults that emerged from lower temperatures (1, 4, and 7 ºC) displayed no significant variation in wing length, there was one distinct trend: wing length was greater in adults that emerged from 4 ºC compared to 1° and 7° C. Similar to previous findings (Marden & Kramer, 1994), increased wing length translated to increased skimming velocity, but skimming speed was not related to incubation temperature. High variation among individuals, coupled with a small sample size, likely obscured any significant effects. *Lednia* often fly just above streams (authors, *pers. obs*.) and, like other small stoneflies (Macneale, Peckarsky, & Likens, 2005), they probably disperse both actively, and passively with strong winds, to nearby streams (Green et al., 2022). This dispersal activity is inferred from their population genetic structure, with *Lednia* populations exhibiting gene flow across neighboring basins (Hotaling et al. 2018). Thus, if longer wings are correlated with better dispersal capacity in *L. tumana*, as they are in other species (Berwaerts, Van Dyck, & Aerts, 2002; Malmqvist, 2000), long-winged individuals should contribute more to population connectivity. Similarly, if short wings are correlated with lower dispersal capacity, increasing stream temperatures may depress connectivity among populations. Given that high-alpine populations of many aquatic insects appear increasingly at risk from climate change, future work should focus on mechanistic links between in-stream conditions and dispersal capabilities.

### The challenge of integrating multiple traits to predict climate vulnerability

Assessing species’ vulnerability to climate change is challenging, especially for species that reside in climatically harsh, remote habitats. Many studies implement a ‘quick and dirty’ approach to vulnerability by measuring critical limits (Huey et al., 2012), which requires the least time and resources. Our study, however, serves as a cautionary tale for this economical approach. In addition to variation in response to high-temperature stress among key traits and life stages as has been shown in other species under thermal stress (Horne, Hirst, Atkinson, Almeda, & Kiørboe, 2019; Kingsolver & Buckley, 2020; Levy et al., 2015), we show that *L. tumana* fails to complete its life cycle at temperatures far below CT_MAX_. Emergence—the moment of transition from an aquatic to a terrestrial phase—is also the most thermally sensitive component of its life history. Rising stream temperatures may therefore imperil populations of *L. tumana* by hindering emergence. This general conclusion is subject to obvious caveats –including lack of information on phenotypic and genotypic variation within and among *L. tumana* populations, the strength, timing, and frequency of selection exerted by high temperatures, and the roles of phenotypic plasticity in promoting persistence in the face of change. In addition, existing or new biotic interactions may also further challenge *Lednia*’s ability to cope with thermal change (Shah et al., 2020b).

## Conclusion

*Lednia tumana* is sensitive to chronic, sub-lethal warming. As climate change proceeds and meltwater sources recede, flows in mountain streams will be reduced (Hotaling et al. 2017). Smaller streams, which contain a majority of freshwater mountain biodiversity, will sustain higher mean temperatures and reach higher maximum temperatures (Birrell et al., 2020). Our results suggest that survival and emergence of *L. tumana* will decline significantly once stream temperatures consistently exceed 7 °C. Over time, low emergence may reduce recruitment, drive population declines, and ultimately lead to population extirpation on local and regional scales. Continued monitoring and physiological testing of populations will help to refine our understanding of climate vulnerability for *L. tumana* and similar taxa worldwide.

## Acknowledgements

This research was supported by a grant from the Rocky Mountain Cooperative Ecosystems Unit (# P17AC00816) and a National Science Foundation postdoctoral fellowship (DBI #1807694) awarded to AAS, and a USGS (#G110-20-W5926) awarded to HAW. *Lednia tumana* specimens were collected under USGS permit (#GLAC-2016-SCI-0007). This work would not have been possible without the help of Brendan Moynahan who liaised between the RMCESU and the University of Montana, October Moynahan who provided excellent logistical and technical support with the incubators, and Rachel Bingham who assisted us in the field and lab. We thank Glacier National Park staff for their logistical support. We also thank Wilco Verberk for providing us with thoughtful comments on an earlier version of this manuscript. Finally, we gratefully acknowledge that our research was conducted on the traditional lands of the Blackfeet, Salish, Kalispel, and Kootenai nations.

