## Supporting Information for "Warming undermines emergence success in a threatened alpine stonefly: a multi-trait perspective on vulnerability to climate change"

1. Wing area measurements

We removed fore- and rear wings from the right side of frozen adults and photographed them under a microscope. Then, using the photographs, we traced a perimeter along each wing with the ‘polygon tool’ in Image J. We calculated the area for each polygon to determine the area of the wing (Fig. S1.).

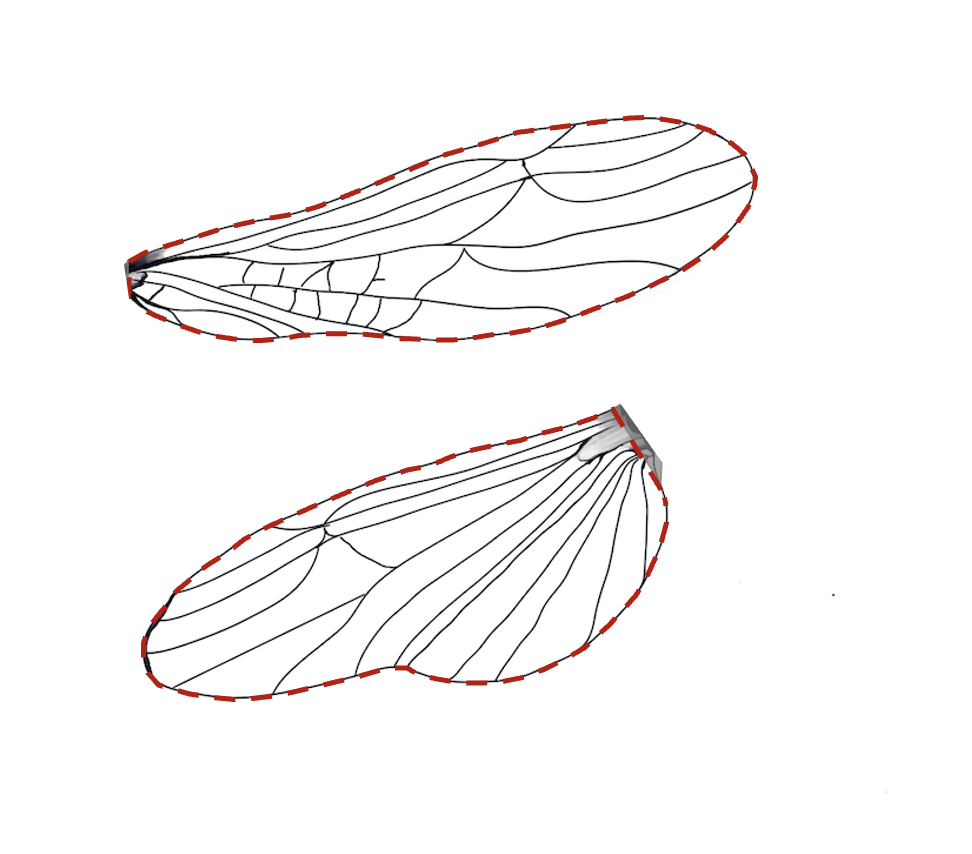

**Figure S1** Schematic demonstrating how wing area measurements were made. The dashed red line represents the polygon trace of a forewing (above) and rear wing (below).

2. Skimming speed and wing length

We analyzed the relationship between skimming speed and wing length for fore- and rear wings. Interestingly, we found a significant relationship between wing length and skimming speed in both types of wings, confirming that wing length is a good predictor of skimming performance. However, possibly due to the very low sample size of emerging adults, we did not find a significant relationship between wing length and incubation temperature.

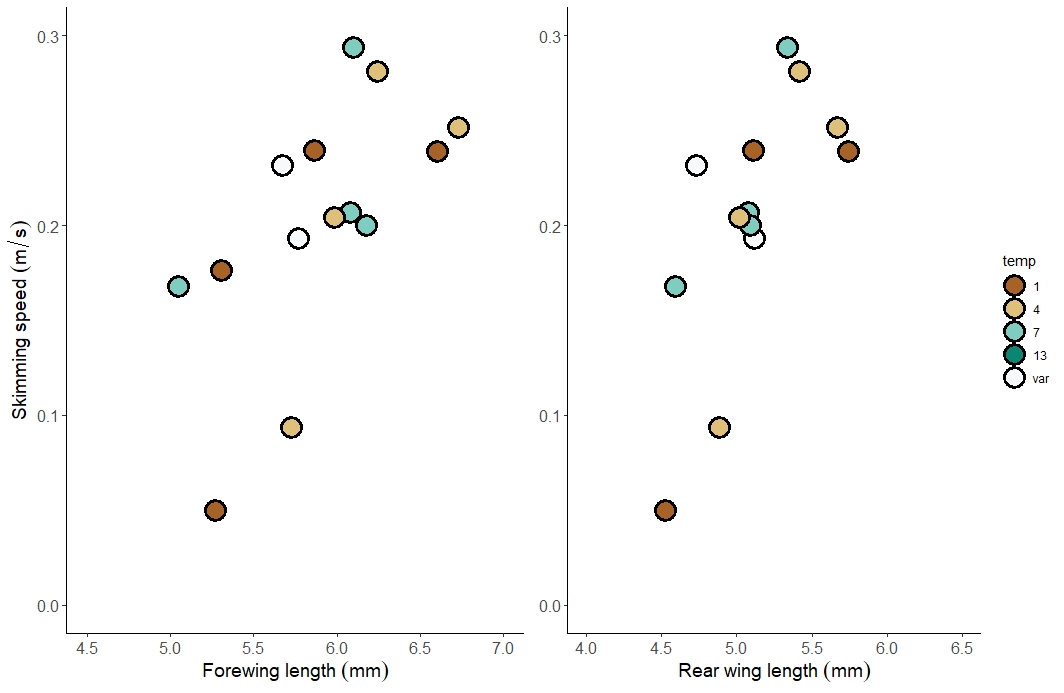

*F*_(1, 12)_ = 8.20, *P* = 0.014, *R^2^* = 0.41

*F*_(1, 11)_ = 9.15, *P* = 0.012, *R^2^* = 0.45

A

B

**Figure S2** Relationship between skimming speed and forewing length (A) and rear wing length (B) for insects reared at different temperatures. For both types of wings, longer wings were strongly associated with higher skimming speeds. Colors indicate incubation temperatures.
